# Evaluation of current and emerging anti-malarial medicines for inhibition of *Toxoplasma gondii* growth in vitro

**DOI:** 10.1101/316455

**Authors:** Joshua B. Radke, Jeremy N. Burrows, Daniel E. Goldberg, L. David Sibley

## Abstract

*Toxoplasma gondii* is a common zoonotic infection of humans and estimates indicate that 1-2 billion people are chronically infected. Although largely asymptomatic, chronic infection poses risk of serious disease due to reactivation should immunity decline. Current therapies for toxoplasmosis only control acute infection caused by actively proliferating tachyzoites but do not eradicate the chronic tissue cyst stages. As well, there are considerable adverse side effects of the most commonly used therapy of combined sulfadiazine and pyrimethamine. Targeting the folate pathway is also an effective treatment for malaria, caused by the related parasites *Plasmodium* spp., suggesting common agents might be used to treat both infections. Here we evaluated currently approved and newly emerging medicines for malaria to determine if such compounds might also prove useful for treating toxoplasmosis. Surprisingly, the majority of anti-malarial compounds being used currently or in development for treatment of malaria were only modestly effective at inhibiting in vitro growth of *T. gondii* tachyzoites. These findings suggest that many essential processes in *P. falciparum* that are targeted by anti-malarial compounds are either divergent, or non-essential in *T. gondii*, thus limiting options for repurposing of current antimalarial medicines for toxoplasmosis.

## Introduction

*Toxoplasma gondii* is a common parasite of animals that causes zoonotic infections in humans. It has diverged from its closest relatives by adopting a broad host range re-enforced by flexible modes of transmission (1). *Toxoplasma gondii* is transmitted by cats, where sexual reproduction in the intestine results in shedding of highly resistant oocysts into the environment (2, 3). Ingestion of oocysts by rodents, and many other intermediate hosts, results in acute infection that is characterized by initial expansion of a fast growing tachyzoite form that disseminates widely throughout the body. Following a vigorous immune response the parasite differentiates into a slow growing, semi-dormant form called the bradyzoite, which inhabits tissue cysts in the muscle and brain (4-6). Human infections are acquired by ingestion of oocysts that contaminate food or water, or by eating undercooked meat that harbors tissue cysts. *Toxoplasma gondii* is a significant cause of serious food borne infection in the USA (7), and it has also been associated with waterborne outbreaks in North and South America (8). Global serological studies indicate that ~25% of humans are chronically infected, although prevalence rates vary widely in different locations (7). Most human infections with *T. gondii* are relatively benign, although they are persistent, as the chronic stages of infection (i.e. bradyzoites within tissue cysts) are not eliminated by the immune response. Additionally, toxoplasmosis can cause serious disease due to congenital infection (9) and in immuncompromised patients (10, 11). Additionally, even healthy adults are at risk due to highly pathogenic strain types that are found some regions such as South America (12, 13).

Toxoplasma is a member of the phylum Apicomplexa, a group of more than 10,000 known species, most of which are parasitic (14). Other apicomplexan parasites of medical importance include *Plasmodium* spp., the causative agent of malaria (15), and *Cryptosporidium* spp., a frequent cause of severe diarrheal disease in young children in developing countries (16). Apicomplexans are most closely related to ciliates and dinoflagellates, but only distantly related to humans, hence they share many key differences from their hosts (17). Although members of the phylum Apicomplexa span 400 mya of evolution (18), they contain many orthologous genes and much of their biology is conserved (19). Among their similar features, *Plasmodium* and *Toxoplasma* contain intact pathways for pyrimidine biosynthesis, while they are purine auxotrophs, and these pathways have been the focus of development of inhibitors to combat both infections (20).

Current therapies for treatment of toxoplasmosis rely primarily on inhibition of the folate pathway in the parasite, although antibiotics developed for treating bacterial infections have also been used with some success (21, 22). The most commonly used treatment is a combination of sulfa drugs with pyrimethamine (i.e. sulfadiazine and pyrimethamine or trimethoprim with sulfamethoxazole). This combination is highly synergistic as the sulfa drug inhibits dihydropteroate synthase (DHPS) while pyrimethamine inhibits dihydrofolate reductase (DHFR), together disrupting tetrahydrofolate levels and blocking DNA synthesis. The combination of pyrimethamine and sulfa drugs is highly effective in blocking replication of tachyzoites but has no activity on bradyzoites within tissue cysts and therefore does not eliminate chronic infection (23). As well, there are significant adverse effects of this treatment regime due to intolerance or allergic reaction to the sulfa component and bone marrow suppression that requires co-administration of leucovorin (24, 25). Additionally, due to potential for inducing developmental defects, this combination is contraindicated during the first two trimesters of pregnancy, but can be effective in reducing clinical severity when given in the third trimester (26). Although alternative therapies such as clindamycin, azithromycin, and other antibiotics, have also been used to treat acute toxoplasmosis, they also do not clear chronic infection (21).

There have been several efforts to identify new drugs for toxoplasmosis based on FDA-approved drugs. Screening of FDA approved drugs has revealed several inhibitors of tachyzoite growth in vitro, most of which were initially developed to treat inflammation (27). Guanabenz, which targets alpha-2 adrenergic receptors and is used for hypertension, additionally shows efficacy in mouse models of toxoplasmosis (28, 29). Finally, treatment of infected cells with tamoxifen, an inhibitor of the estrogen receptor, leads to parasite clearance due to an autophagy-related process (30). Although such compounds provide promising leads, they do not allow selective inhibition of the parasite since they were originally optimized to modulate host processes. Hence, there is a need for new treatments that are more selective, less toxic, and effective at eliminating chronic infection by *T. gondii*.

One potential source for new drugs to treat toxoplasmosis would be repurposing of medicines that have been developed for malaria, a concept that is based on their shared ancestry and similar biology. One example is the use of pyrimethamine-sulfa drug combinations to treat toxoplasmosis. Similarly, Fansidar (sulfadoxine and pyrimethamine) was historically effective against *P. falciparum*. However, due to the global spread of anti-folate (31) and chloroquine resistance (32), the first line of treatment for malaria has shifted to the natural product artemisinin, which is a sequiterpene lactone that contains an endoperoxide bridge that is key to its activity (33). A number of semi-synthetic variants have been produced including artesunate, artemether, and artemisone, which are more soluble prodrug forms that are rapidly converted to dihydroartemisinin in plasma (31). Artemisinin derivatives also inhibit replication of *T. gondii* in vitro (34-36) and are partially effective in murine infection models, although they do not eliminate chronic infection (37, 38).

Considerable effort has been expended to develop new generation anti-malarials based on large-scale phenotypic high throughout screens (HTS) for inhibition of asexual blood stage forms of *P. falciparum* (39). A number of the resulting hits were prioritized by the Medicines for Malaria Venture (MMV) based on chemical properties as well as activity to define a core set of compounds for inclusion in the Malaria Box and Pathogen Box projects (https://www.pathogenbox.org/). Combined with genomic analyses of evolved resistant mutants, these screening efforts have led to identification of new leads that target essential steps in the parasite (39). One of the first new active malarial compounds to be identified by a screening/genomics approach was the class of compounds known as spiroindolones (40), including the analog KAE609 that proved effective in curing mice of *P. berghei* infection with a single oral dose (41). Whole genome sequencing of resistant mutants, and subsequent genetic confirmation, indicated that spiroindolones target the cation transporter PfATP4 (42), thereby disrupting sodium transport in the parasite (43). A similar strategy of whole genome sequencing of resistant mutants has led to the identification of several tRNA synthases as targets of potent antimalarial candidate compounds and mutations in the PfCarl (cyclic amine resistance locus) gene that medicates resistance to potent imidazolopiperizines (39).

The availability of the Malaria and Pathogen Box (https://www.pathogenbox.org/) collections has made it possible to expand the analysis of these compounds to other pathogens (44). Analysis of compounds in the Malaria Box for inhibition of in vitro growth of *T. gondii* identified seven compounds with EC_50_ ≤ 5 μM including a piperazine acetamide with an EC_50_ < 0.19 uM (45). The hit rate of ~ 2% observed in this study is higher than typically seen in typical HTS; however, it might be considered low based on the premise that *T. gondii* and *P. falciparum* are members of the same phylum and share much of their underlying biology. Several of the active compounds contain a quinolone moiety, suggesting they may be active due to their resemblance to endochin-like quinolones (46) and atovaquone (47), which act on the bc1 complex and that are active against *T. gondii*. A second study that also employed the Malaria Box reported a much higher hit rate, with 49 compounds out of ~400 showing EC_50_ values ≤ 1 μM when tested for inhibition of tachyzoite growth in vitro (48); the difference in hit rate being attributed to methodology. Although a number of these compounds were also inhibitors of *P. falciparum*, the overall correlation in potency between these two parasites was low (48). A similar screen for tachyzoite in vitro growth inhibition by compounds in the Pathogen Box identified four compounds with EC_50_ values ≤ 1.0 μM and selectivity indices of > 4 (49). Among the more potent compounds identified was buparvaquone, which is a naphthoquinone that also inhibits mitochondrial electron transport. This study also reported that many compounds active against *Plasmodium* did not show appreciable inhibition of *T. gondii*.

To complement previous efforts that have focused on the early preclinical leads found in the Malaria and Pathogen Box collections, we focused here on approved medicines for malaria, new anti-malarial candidates or emerging leads that are in the global malaria portfolio, many of which with Medicines for Malaria Venture (MMV) (39, 50). Many of these compounds show excellent potential for treatment of malaria, and have advanced through a number of preclinical safety checks and in some cases clinical studies, and include a number of currently used medicines. The rationale for this project was that if active compounds were found among this set, they might be readily repurposed for treatment of toxoplasmosis or give information on biological pathways to target in *T. gondii*.

## Results and Discussion

We tested 81 compounds including a number of current medicines used for treatment of malaria and candidates that are in late preclinical development or undergoing current clinical trials. Although some individual compounds had been tested on *T. gondii* previously, many are new compounds and this represents the first time this set of compounds has been compared side by side in the same assay. We tested them in parallel using a multi-well plate assay that monitors in vitro growth of *T. gondii* tachyzoites based on firefly luciferase expression. Initially, we evaluated each of the compounds at a single concentration (10 μM) in triplicate assays to define those that showed > 50 % growth inhibition. Based on this cutoff, 52 compounds were chosen for further analysis based on duplicate 10-point dilution series that were used to define the EC_50_ values for growth inhibition. Results of the screen are summarized in Table 1 and Figure 1, where compounds are ranked by relative potency.

**Table 1.** Sumary of EC_50_ values for inhibtion of *T. gondii* growth in vitro.

| Compound Name | Chemical class | Known or suspected mechanism of action (resistance mechanism) | EC<sub>50</sub> log inhib vs normalized response (μM) |
| --- | --- | --- | --- |
| Clindamycin | macrolide - lincosamide | Protein synthesis - bacterial 50S ribosomal subunit | 0.0081 |
| Trans-Mirincamycin | macrolide - lincosamide | Protein synthesis - bacterial 50S ribosomal subunit | 0.0140 |
| Cyclohexamide | glutarimide | Protein synthesis - eukaryotic - elongation | 0.0161 |
| Cis-Mirincamycin | macrolide - lincosamide | Protein synthesis - bacterial 50S ribosomal subunit | 0.0197 |
| ELQ-300 | quinolone | Mitochondrial bc1 complex- Qi | 0.0577 |
| Artemisone | sesquiterpene lactone | peroxide-mediated, oxidative damage (Kelch 13) | 0.0755 |
| Azithromycin | macrolide - azalide | Protein synthesis - bacterial 50S ribosomal subunit | 0.1654 |
| Atovaquone | hydroxynaphthylquinone | Mitochondrial bc1 complex- Qo | 0.2441 |
| Pyrimethamine | pyrimidine derivative | DHFR | 0.2538 |
| KAE609 | spirotetrahydro β-carboline | ATP4 | 0.2561 |
| Artemether | sesquiterpene lactone | peroxide-mediated, oxidative damage (Kelch 13) | 0.2861 |
| Methylene blue | phenothiazin | Uncertain | 0.2958 |
| Artesunate | sesquiterpene lactone | peroxide-mediated, oxidative damage (Kelch 13) | 0.5989 |
| Ro 47-7737 | 4-aminoquinoline (bis) | Hemozoin formation | 0.6508 |
| Artemisinin | sesquiterpene lactone | peroxide-mediated, oxidative damage (Kelch 13) | 0.7595 |
| BIX-01294 | diaminoquinazoline | Histone methyl transferase | 0.8792 |
| Phenylequine | 4-aminoquinoline | Hemozoin formation | 0.9362 |
| MMV688558 | pantothenamide | Co-enzyme A | 0.9526 |
| Doxycycline | macrolide - tetracycline | Protein synthesis - bacterial | 1.0080 |
| Thiostrepton | cyclic oligopeptide - thiopeptide | Protein synthesis - bacterial 50S ribosomal subunit | 1.1310 |
| Dihydroartemisinin | sesquiterpene lactone | peroxide-mediated, oxidative damage (Kelch 13) | 1.1390 |
| Pyronaridine | 4-anilino-quinoline | Hemozoin formation and novel | 1.2430 |
| KAF246 | spirotetrahydro β-carboline | ATP4 | 1.4000 |
| Chlorproguanil | biguanide - prodrug | DHFR after cyclization - and unique | 1.4330 |
| 2k | 4-anilino-quinoline | Hemozoin formation | 1.4560 |
| NPC-1161B | 8-aminoquinoline | Uncertain | 1.4680 |
| Mefloquine (racemic) | quinoline amino-alcohol | Uncertain (Pfmdr1) | 1.4900 |
| Sitamaquine | 8-aminoquinoline | Uncertain | 1.5800 |
| Cladosporin | isocoumarin | Lysyl t-RNA synthase | 1.5820 |
| (+)-Mefloquine | quinoline amino-alcohol | Uncertain (Pfmdr1) | 1.661 |
| AQ-13 | 4-aminoquinoline | Hemozoin formation | 1.6460 |
| AZ412 | triaminopyrimidine | Vacuolar ATPase synthase / V-type H<sup>+</sup> ATPase | 1.7560 |
| Pamaquine | 8-aminoquinoline | Uncertain | 1.7840 |
| Halofantrine | amino alcohol | Hemozoin formation | 1.8140 |
| OZ277 | trioxolane - synthetic endo-peroxide | peroxide-mediated, oxidative damage | 1.8570 |
| N-desethyl amodiaquine | 4-anilino-quinoline | Hemozoin formation | 1.8670 |
| Dapsone | sulfone | Dihydropteroate synthesis | 1.9430 |
| Tafenoquine | 8-aminoquinoline | Uncertain | 2.1160 |
| Amodiaquine | 4-anilino-quinoline | Hemozoin formation | 2.2260 |
| AN13762 | oxaborole | Uncertain | 2.7030 |
| OZ439 | trioxolane - synthetic endo-peroxide | peroxide-mediated, oxidative damage | 2.7220 |
| Cycloguanil | cyclic-biguanide | DHFR | 2.7580 |
| UCT944 | aminopyrazine | PI4K | 2.7900 |
| KAF156 | imidazolopiperazine | (PfCarl) | 3.0620 |
| Sulfamethoxazole | sulfonamide | Dihydropteroate synthesis | 3.1530 |
| MK-4815 | aminocresol | Hemozoin formation | 3.3010 |
| UCT048 | aminopyridine | PI4K | 3.4340 |
| Primaquine | 8-aminoquinoline | Uncertain | 3.5140 |
| Pentamidine | bisamidine derivative | Uncertain | 3.6790 |
| Proguanil | biguanide - prodrug | DHFR after cyclization - and unique | 4.4700 |
| Sulfadiazine | sulfonamide | Dihydropteroate synthesis | 4.5220 |
| 21A092 | pyrazoleamide | ATP4 | 4.6370 |

**Figure 1.**
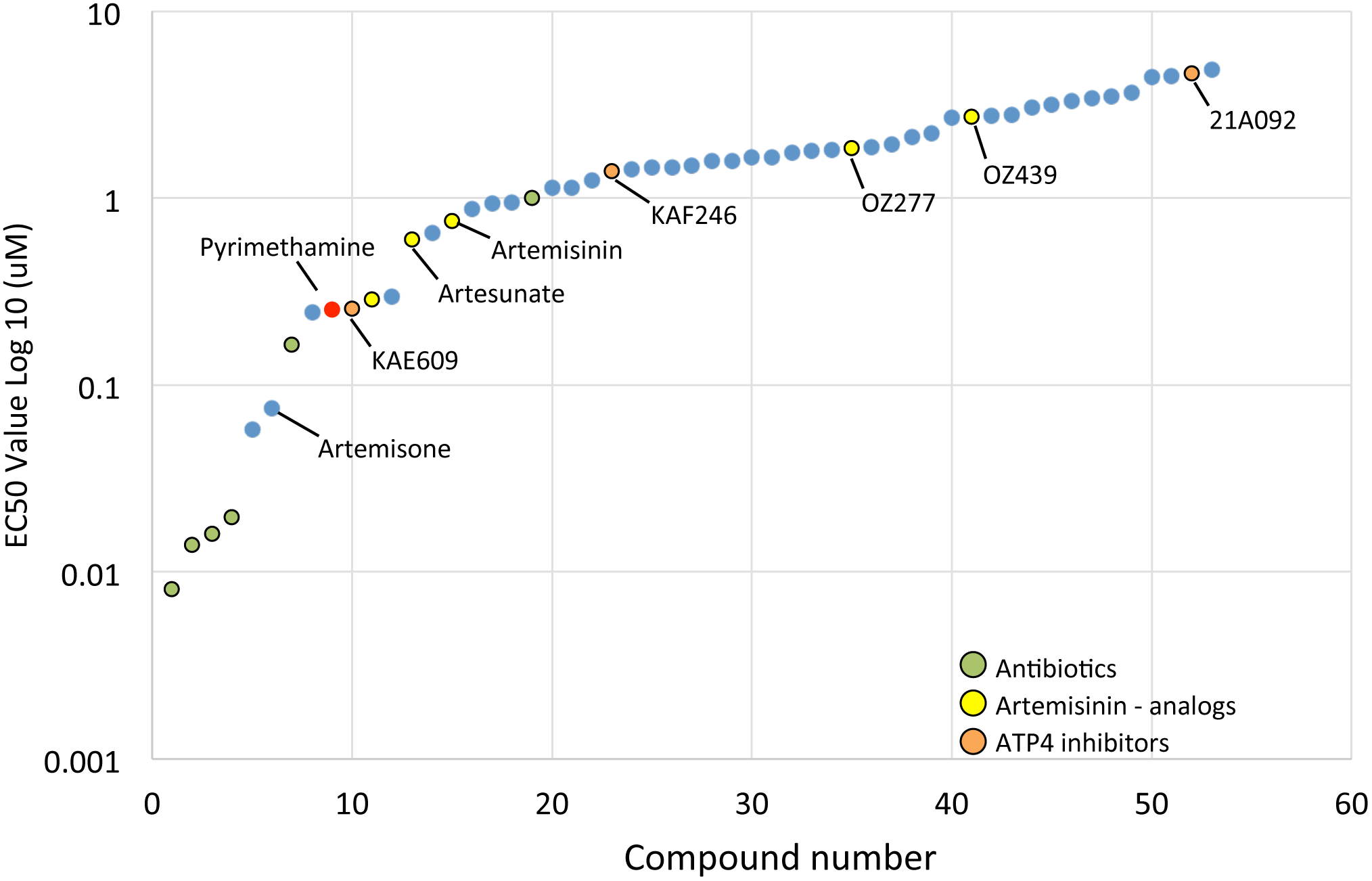
Rank order of compounds in terms of EC_50_ values for inhibition of *T. gondii* tachyzoite growth in vitro. Values represent mean of two independent 10-point titrations that were used to derive EC_50_ values by doseresponse curve fits (see methods). Compounds of interest are highlighted including pyrimethamine (red dot), artemisinin and related compounds (yellow), inhibitors of PfATP4 (orange), and antibiotics (green). See Table 1 for complete EC_50_ values.

Among the set of 52 compounds, 18 of them showed EC_50_ values of < 1 μM, including pyrimethamine (Figure 1, red dot), consistent with previously reported values for the activity of this DHFR inhibitor on growth of *T. gondii* in vitro (51, 52). Other inhibitors of DHFR were considerably less potent, including methotrexate (53, 54) and cycloguanil (55, 56), consistent with previous reports (Table 1). As expected, sulfa drugs were also less potent including sulfadiazine, sulfamethoxoazole, and dapsone (Table 1), consistent with previous reports of their in vitro activity (52, 57, 58). The low activity of these molecules vitro may reflect high levels of *p*-araminobenzoic acid in culture medium, as this metabolite acts competitively with these inhibitors of DHPS. Although sulfa drugs are not effective when used alone, as part of the current combination therapy they are highly synergistic with pyrimethamine (52, 59). Among the potent compounds that have not been reported previously was methylene blue, a phenothiazin dye that is being evaluated as a transmission blocking compound for malaria (50). Also included among the most actives were several antibiotics that target prokaryotic protein synthesis by disrupting ribosomes including the lincosamides clindamycin and mirincamycin, and the macrolide azithromycin (Table 1). Interesting, although doxycycline was active in inhibiting *T. gondii* growth, tetracycline showed almost no activity, consistent with a previous report (60). A number of antibiotics have previously been shown to be active on *T. gondii* (59, 61-64) and their mechanism of action is likely due to inhibition of protein synthesis in the apicoplast (65). The main limitation to use of broad-spectrum antibiotics for treatment of toxoplasmosis is their potent activity on the endogenous bacterial flora in the microbiome leading to disbiosis and gastrointestinal distress, thus increasing the risk of *C. difficile* infection (66). The use of such agents that target bacteria also increases the risk of unwanted emergence of resistance among other classes of pathogens.

Other potent inhibitors include the endochin-like quinolone ELQ-300 that targets the Qi site in the cytochrome bc1 complex (67) and atovaquone that targets the Qo site (68) in the cytochrome bc1 complex of the mitochondrial respiratory chain (Table 1). Previous studies have shown that atovaquone is effective in blocking *T. gondii* replication in vitro and in reducing cyst numbers in chronically infected mice (52, 62), as well as suppressing reactivation of chronic infection in an immunocomprmised mouse model (69). However, prior experience with atovoquone in humans with toxoplasmosis includes several examples of therapeutic failure (47, 70), possibly due to resistance arising, although the mechanism was not confirmed at the molecular level. Similar to atovoquone, ELQ-300 is potent in inhibiting parasite growth in vitro and in reducing cyst numbers in the brains of chronically infected mice (46). The main issue with ELQ-300 (46), and related quinolone compounds (71), is their low solubility that reduces oral bioavailability. Consequently, efficacy trials in murine models of toxoplasmosis have relied on parenteral administration of the compounds. This limitation has been partially mitigated by production of esterified pro-drugs that get activated in vitro, allowing for oral treatment that was protective in a mouse model for *P. yoelli* (72). Given that multiple quinolone containing compounds that affect mitochondrial electron transport are active against *T. gondii*, including against bradyzoites in tissue cysts, this pathway remains an important target for further investigation.

Drugs that have traditionally been used to treat malaria were much less potent in inhibiting *T. gondii* including both 4-amino and 8-amino quinolines (Table 1). Chloroquine and a variety of related 4-aminoquniolines are active against asexual parasite stages of *Plasmodium* that replicate in red blood cells, where these compounds are thought to inhibit hemozoin formation within the parasite’s digestive food vacuole (73). Among this class of compounds, bisquinoline and benzylquine were the most active with EC_50_ values of less than 1 μM, while other derivatives were less potent against *T. gondii* (Table 1). These compounds are thought to target hemozoin formation during hemoglobin degradation in *Plasmodium* (73), and the lack of an analogous digestive pathway in *T. gondii* may explain the lack of potency of most members of this class. However, the fact that several 4-amino quinolones and 4-anilino compounds (i.e. pyronaridine, amodiaquine) showed modest potency in inhibiting *T. gondii*, suggests that they target another important process (Table 1).

A number of 8-aminoquinolones are also effective for treatment of malaria, although their mechanism of action remains uncertain. Primaquine has traditionally been used to treat the dormant hypnozoite stage of *P. vivax* and it is also effective against *P. falciparum* gametocytes. The main deficiency of this compound is its toxicity in patients with G6PD deficiency. A number of 8-aminoquinoline derivatives, such as tafenoquine, lack some of the undesirable effects of primaquine and are also being advanced for preventing relapse of *P. vivax* (74). Unfortunately, the 8-aminoquinolines as a class were largely inactive against *T. gondii* tachyzoites (Table 1). However, given their differential activity on semi-dormant stages of *Plasmodium* development (i.e. gametocytes and hypnozoites) it would be interesting to test these compounds on bradyzoites of *T. gondii*.

Artemisinin derivatives have become the mainstay of combined therapy against severe and uncomplicated malaria (31). Artemsinin is potent across the stages of intraerythrocytic development and this activity has been attributed to hemoglobin degradation and release of free heme, which is thought to activate the endoperoxide bridge, likely forming adducts with multiple targets (33). More recent efforts have focused on completely synthetic endoperoxides, some of which show greater metabolic stability in vivo, and which could reduce reliance on the natural product produced from *Artemesia* cultivation (75). Consistent with prior studies (35, 37), a number of artemisinin derivatives were modestly active in inhibiting *T. gondii* growth (Table 1, Figure 2). Artemisone and artemether were among the most active derivatives, while deoxyartemesinin was inactive, indicating that activity is dependent on the endoperoxide moiety. Unfortunately, more stable trioxolane synthetic peroxides such as OZ439 and OZ277 were much less active on *T. gondii* (Table 1, Figure 2). Even the most potent artemisinin derivatives are several orders of magnitude less effective against *T. gondii* compared to *P. falciparum*, thus limiting their potential as therapeutic options for toxoplasmosis.

**Figure 2.**
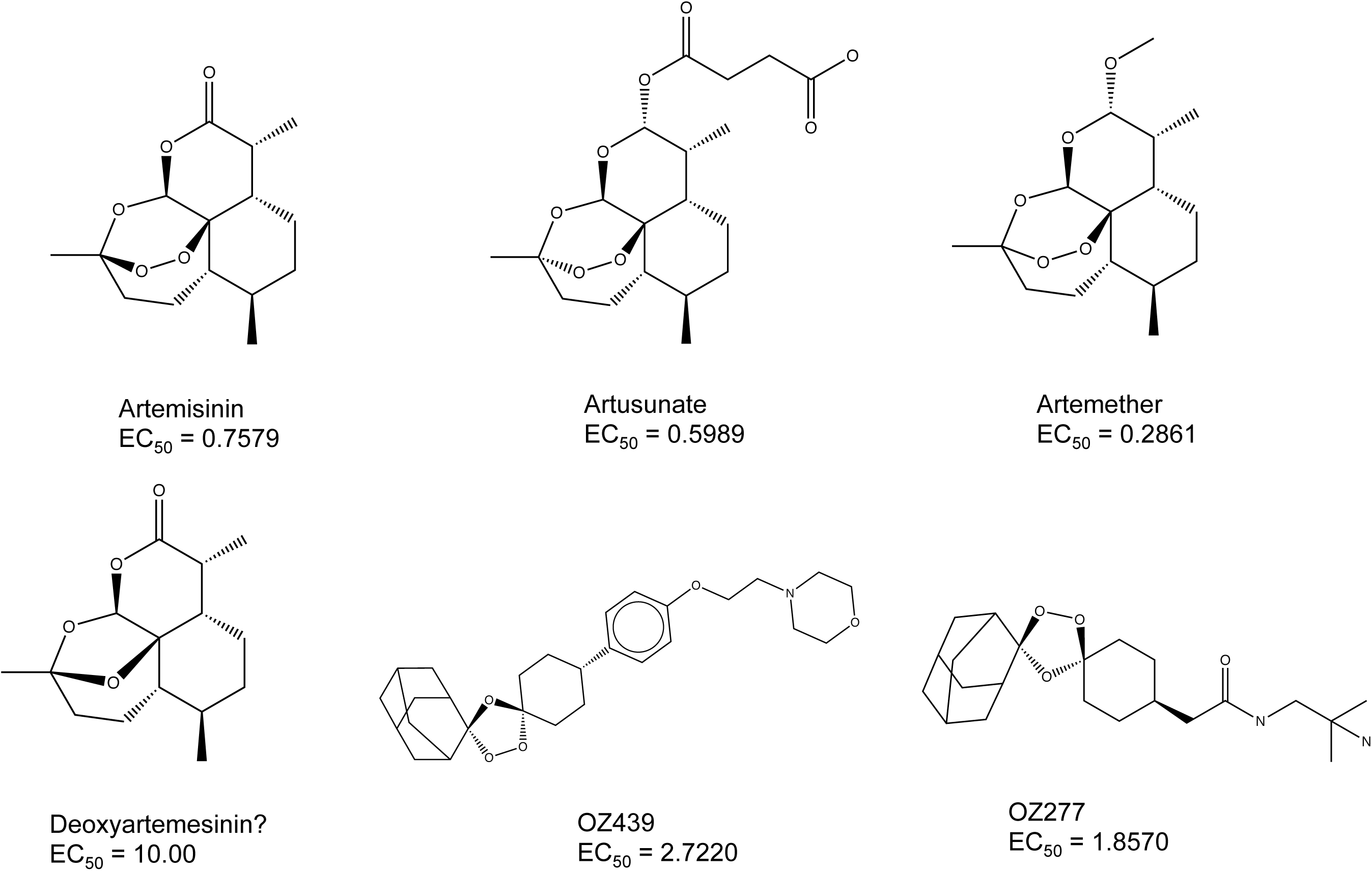
Structures and EC_50_ values for artemesinin and related analogs artusunate and artemether. Deoxyartemesinin, which is inactive, is shown for comparison. Several synthetic trioxanes are also illustrated. See Table 1 for complete EC_50_ values.

Several different chemical scaffolds have been shown to inhibit the P-type cation translocating ATPase in *P. falciparum* known as PfATP4, which resides in the parasite plasma membrane where it modulates cytosolic sodium levels by active extrusion of Na^+^ in exchange for H^+^ (76). Among the most potent PfATP4 inhibitors are the spiroindolones that are currently in clinical trials for treatment of infection by *P. falciparum* (50). The spiroindolone KAE609 (also known as NITD609), which is highly active on *P. falciparum* (41-43), was among the more active molecules studied here for inhibition of *T. gondii* growth (EC_50_ ~ 0.25 μM) (Table 1, Figure 3). KAE609 was substantially more potent than a second spiroindolone analog KAF246 (EC_50_ = 1.4 μM) (Table 1, Figure 3), although these two compounds differ only by substitution of Cl for F on the indole group (Figure 3). KAE609 was previously reported to inhibit *T. gondii* growth in vitro with a 50% decrease in parasite growth at 1 μM (reported as MIC_50_) and in a mouse model for acute toxoplasmosis when the compound was administered at 100 mg/kg (given orally on the day of infection and the day after infection) (77). Compound 21A092, which belongs to a different scaffold known as a pyrazoleamide, also targets PfATP4 and is highly active on *P. falciparum* (78), but was much less potent on *T. gondii* (Table 1). Unfortunately, even potent spiroindoles like KAE609 show much greater potency on *P. falciparum* (EC_50_ ~1 nM) (42) than *T. gondii*, despite the fact that the proposed binding site in PfATP4 is highly conserved, including sites that result in resistance when mutated (77). Interestingly, the relative potency between the ATP4 inhibitors (highest to lowest) KAE609, PA92 and SJ733 on *P. falciparum* appears to match that in *T. gondii*. A number of other chemical scaffolds have also been shown to affect PfATP4 in *P. falciparum* and a previous screen of the Malaria Box identified a number of compounds that likely target PfATP4 (76) as inhibitors of *T. gondii* (48). As many analogs of the spiroindolones, and other scaffolds that act on PfATP4, are available, it may be worth further investigation of this target to identify more potent inhibitors of *T. gondii*.

**Figure 3.**
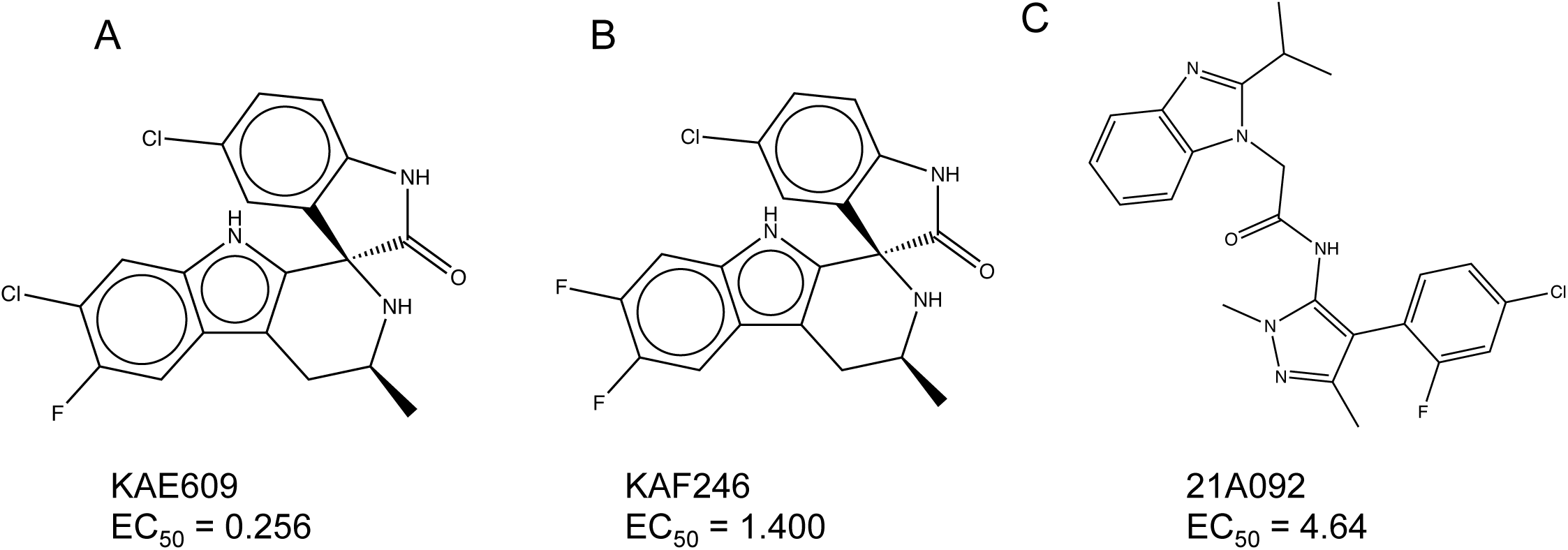
Structures and EC_50_ values for several PfATP4 inhibitors including two spiroindolones (KAE609 and KAF246) and a structurally unrelated pyrazoleamide (21A092). See Table 1 for complete EC_50_ values.

Additionally, potent new antimalarial compounds that have been recently identified showed only modest activity against *T. gondii* with EC_50_ values that ranged from 1 – 5 μM (Table 1). Included in this group was KAF156, an imidazolopiperazine that showed promising results in a recent clinical trial of *P. falciparum* malaria (79). Additionally, several inhibitors of phosphoinositol 4 kinase (PI4K) showed modest (i.e. UTC944), or no appreciable (i.e. KDU691), activity against *T. gondii* growth, despite having excellent potency against *P. falciparum* (80). Similarly, cladosporin, which targets lysyl tRNA synthase in *P. falciparum* (81, 82), showed only modest activity against *T. gondii* (Table 1). The reasons for the much lower activities observed on *T. gondii* is uncertain but it may reflect differences in the molecular targets of these compounds or differences in the extent with which these targets play essential roles in the biology of *T. gondii* vs. *P. falciparum*. Alternatively, these differences might arise from differences in the intracellular compartment that affect access of the compounds or from a greater number of efflux mechanisms in *T. gondii*. Regardless of the precise reasons, these differences in sensitivity provide a rationale to explore a more diverse collection of compounds than studied here, with the potential that other analogs within these chemical scaffolds will be found to be more effective on *T. gondii*.

Targeting conserved and essential pathways may thus offer greater advantage for finding compounds with a broader spectrum of activity. One potential example is pyrimidine biosynthesis that is conserved in both *T. gondii* and *P. falciparum*. In particular, *Plasmodium* lacks pyrimidine salvage enzymes and thus is reliant on biosynthesis for RNA and DNA synthesis. Targeted screens have advanced new triazolopyrimidine compounds as inhibitors of *P. falciparum* dihydroorotate dehydrogenase (DHODH), thus blocking pyrimidine biosynthesis (83). One such analog DSM265 is active against both liver and blood stages of *P. falciparum*, shows excellent pharmacokinetic and safety properties, and has advanced to clinical trials for *P. falciparum* malaria (84). DHODH is also essential in for pyrimidine biosynthesis in *T. gondii*, as well as performing another essential function in mitochondria (85). However, DSM265 was not effective at inhibiting *T. gondii* growth in vitro (Table 1). This difference may reflect that fact that DSM265 has been carefully selected for potency on *P. falciparum* hence this may reflect a difference in the molecular target, suggesting that other analogs may be more effective. Alternatively this result that may be due to the capacity of *T. gondii* to salvage uracil (86). Hence, even apparently conserved pathways present challenges for identification of potent inhibitors both due to potential molecular differences in the target and/or alternative metabolic routes in these two parasites.

## Conclusions

We evaluated 80 compounds that are used as current therapy for malaria, or which are in late stage development, for their ability to inhibit the growth of the related apicomplexan parasite *T. gondii*. The most active compounds identified were previously known agents including lincosamide and macrolide antibiotics that target the apicoplast and quinolones that target the bc1 complex in the mitochondria. Consistent with this pattern, artemisinin and related analogs, were modestly potent, while several new generation trioxanes showed very little activity. Moreover, traditional drugs used against malaria including 4-amino and 8-aminoquinolines showed very little activity against *T. gondii*. Similarly, a number of newly identified compound classes that target novel pathways in *Plasmodium* showed limited activity against *T. gondii* including spiroindolones, which inhibit PfATP4, as well as compounds that target PI4K, lysyl tRNA synthase, and others. These findings may suggest that current malaria drugs target pathways that are not conserved in these two parasites, or alternatively that differences in the molecular target will require different analogs to effectively target each of these parasites. Hence, identifying new treatments for toxoplasmosis will require a concerted effort to identify potent inhibitors of essential targets in this organism.

## Materials and Methods

### Cell culture and parasite propagation

Tachyzoites of ME49 strain encoding a transgenic copy of firefly luciferase (type II, ME49-FLuc) (87) were continually passaged in confluent monolayers of human foreskin fibroblasts (HFF) cultured in DMEM supplemented with 10% fetal bovine serum (FBS), glutamine (10 mM) and gentamycin (10 μg/mL). To isolate parasites, heavily infected cultures of late-stage vacuoles containing replicating tachyzoites were scraped, force lysed through a 23g needle and residual host cell material removed using a polycarbonate filter (3 micron pore). The parasites were then counted, diluted in fresh culture medium and added to 96-well plates as described below. All HFF and parasite cultures were grown in a 37°C incubator supplemented with 5% CO_2_ and were verified to be mycoplasma free using the e-Myco Plus kit (Intron Biotechnology).

### Luciferase based growth inhibition assays

HFF cells were plated in white, clear-bottom 96-well plates (Costar #3610) and incubated to confluency. Only the inner 60 wells were used to reduce variability due to edge affect with outside wells. Compounds KAF246 and KDU691 were obtained from Novartis Tropical Research Institute. The remaining compounds were provided by MMV. All compounds were prepared as 10 mM stock in 100% DMSO and stored at -80°C until use. After dilution, the final DMSO concentration for all experimental wells was 0.1%. Pyrimethamine (Sigma-Aldrich, #46706) was reconstituted in 100% DMSO at 5 mM stock concentration and stored at -20°C until use.

For the single point assay, 5x10^3^ ME49-FLuc parasites (in 100 μL volume) were inoculated into plates containing 100 μL of 2x compound (10 μM final concentration, 200 μL final well volume), incubated for 72h at 37°C and luciferase activity evaluated using the Luciferase Assay System (Promega, E1501) according manufacturers protocol. Briefly, culture medium was aspirated and replaced with 40 μL of 1x Passive Lysis buffer (1xPLB, Promega, E1531) and incubated for 5 min at room temperature (RT). Luciferase activity was measured on a Beckman Coulter integrated and automated platform using the following protocol: Inject 100 μL of Luciferase Assay Reagent (LAR), shake 1 sec and read 10 sec post-injection. Only compounds that showed greater than 50% growth inhibition over DMSO control were selected for EC_50_ determination (average of 3 biological replicates). Liquid handling steps (media exchange, compound dilution and addition and luciferase assay steps) were performed on a Biomek Dual Pod FX system using the SAMI EX software as part of the High Throughput Screening Center at Washington University School of Medicine.

For compounds demonstrated >50% growth inhibition at 10 μM, EC_50_ values were determined form a 10-point dose-response curve. Briefly, 5x10^3 ME49-FLuc parasites (100 μL/well) were added to a 96-well plate that contained 100 uL of 2x compound (1x final compound concentration, 200 μL total well volume,) and allowed to replicate for 72h prior to preparation for luciferase assay. All experimental steps, growth conditions and luciferase assay protocols were completed as described above. Compounds were tested using a 3-fold dilution series from 10 μM to 0.001 μM with all wells containing a final concentration of 0.1% DMSO. Pyrimethamine (2.5 μM, positive control) and DMSO (vehicle) controls were added to the outside wells of all plates as controls. The EC_50_ data are presented as the average of two biological replicates.

### Statistics

All results are presented as the average of two or more biological replicates. Linear regression analysis and dose-response inhibition (Log (inhibitor) vs. normalized response – variable slope) or (Log (inhibitor) vs. normalized response) were performed in Prism 7 (GraphPad Software, Inc.).

## Acknowledgement

We thank Dr. Thierry Diagana, Novartis Institute for Tropical Diseases, for providing KAF246 (aka NITD246) and KDU691, Jennifer Barks for assistance with cell culture, and Dr. Maxine Ilagan, High-Throughput Screening Center at Washington University School of Medicine, for assistance with compound screening. Supported by the National Institute of Allergy and Infectious Diseases of the National Institutes of Health under Award Number U19AI109725.

## Ancillary Information

### Supporting Information

### Author contributions

JBR designed or performed screening assays; JBR and LDS analyzed primary screening data and generated figures; JB and DEG reviewed the findings and provided scientific advice; JBR and LDS wrote the manuscript with input from all authors.

## Abbreviations used

ART: artemisinin
DHFR: dihydrofolate reductase
DHODH: dihydroorotate dehydrogenase
PfATP4: adenosine triphophosphatase 4
HFF: human foreskin fibroblasts
HTS: high throughput screen
PI4K: phosphotidylinositol 4 phosphate kinase
EC: effective concentration

## References

1. Lorenzi H, Khan A, Behnke MS, Namasivayam S, Swapna LS, Hadjithomas M, Karamycheva S, Pinney D, Brunk BP, Ajioka JW, Ajzenberg D, Boothroyd JC, Boyle JP, Darde ML, Diaz-Miranda MA, Dubey JP, Fritz HM, Gennari SM, Gregory BD, Kim K, Saeij JP, Su C, White MW, Zhu XQ, Howe DK, Rosenthal BM, Grigg ME, Parkinson J, Liu L, Kissinger JC, Roos DS, Sibley LD. Local admixture of amplified and diversified secreted pathogenesis determinants shapes mosaic Toxoplasma gondii genomes. Nature communications. 2016;7:10147. PMID: 26738725; 4729833.

2. Dubey JP, Miller NL, Frenkel JK. The *Toxoplasma gondii* oocyst from cat feces. Journal of Experimental Medicine. 1970;132(4):636–662.

3. Frenkel JK, Dubey JP, Miller NL. *Toxoplasma gondii*: fecal forms separated from eggs of the nematode toxocara cati. Science. 1969;164:432–433.

4. Dubey JP. Bradyzoite-induced murine toxoplasmosis: stage conversion pathogenesis, and tissue cyst formation in mice fed bradyzoites of different strains of *Toxoplasma gondii*. J Euk Microb. 1997;44(6):592–602.

5. Dubey JP, Frenkel JK. Experimental *toxoplasma* infection in mice with strains producing oocysts. Journal of Parasitology. 1973;59(3):505–512.

6. Dubey JP, Speer CA, Shen SK, Kwok OCH, Blixt JA. Oocyst-induced murine Toxoplasmosis: life cycle, pathogenicity, and stage conversion in mice fed *Toxoplasma gondii* oocysts. Journal of Parasitology. 1997;83:870–882.

7. Pappas G, Roussos N, Falagas ME. Toxoplasmosis snapshots: global status of *Toxoplasma gondii* seroprevalence and implications for pregnancy and congenital toxoplasmosis. Int J Parasitol. 2009;39(12):1385–1394. PMID: 19433092.

8. Jones JL, Dubey JP. Waterborne toxoplasmosis--recent developments. Exp Parasitol. 2010;124(1):10–25. PMID: 19324041.

9. Torgerson PR, Mastroiacovo P. The global burden of congenital toxoplasmosis: a systematic review. Bull World Health Organ. 2013;91(7):501–508. PMID: 23825877; 3699792.

10. Luft BJ, Remington JS. Toxoplasmic encephalitis in AIDS. Clin Infect Dis. 1992;15:211–222.

11. Israelski DM, Remington JS. Toxoplasmosis in the non-AIDS immunocompromised host. Curr Clin Top Infect Dis. 1993;13:322–356.

12. Pfaff AW, de-la-Torre A, Rochet E, Brunet J, Sabou M, Sauer A, Bourcier T, Gomez-Marin JE, Candolfi E. New clinical and experimental insights into Old World and neotropical ocular toxoplasmosis. Int J Parasitol. 2013. PMID: 24200675.

13. Demar M, Hommel D, Djossou F, Peneau C, Boukhari R, Louvel D, Bourbigot AM, Nasser V, Ajzenberg D, Darde ML, Carme B. Acute toxoplasmoses in immunocompetent patients hospitalized in an intensive care unit in French Guiana. Clin Microbiol Infect. 2012;18(7):E221–231. PMID: 21958195.

14. Pawlowski J, Audic S, Adl S, Bass D, Belbahri L, Berney C, Bowser SS, Cepicka I, Decelle J, Dunthorn M, Fiore-Donno AM, Gile GH, Holzmann M, Jahn R, Jirku M, Keeling PJ, Kostka M, Kudryavtsev A, Lara E, Lukes J, Mann DG, Mitchell EA, Nitsche F, Romeralo M, Saunders GW, Simpson AG, Smirnov AV, Spouge JL, Stern RF, Stoeck T, Zimmermann J, Schindel D, de Vargas C. CBOL protist working group: barcoding eukaryotic richness beyond the animal, plant, and fungal kingdoms. PLoS biology. 2012;10(11):e1001419. PMID: 23139639; 3491025.

15. Miller LH, Ackerman HC, Su XZ, Wellems TE. Malaria biology and disease pathogenesis: insights for new treatments. Nat Med. 2013;19(2):156–167. PMID: 23389616.

16. Kotloff KL, Nataro JP, Blackwelder WC, Nasrin D, Farag TH, Panchalingam S, Wu Y, Sow SO, Sur D, Breiman RF, Faruque AS, Zaidi AK, Saha D, Alonso PL, Tamboura B, Sanogo D, Onwuchekwa U, Manna B, Ramamurthy T, Kanungo S, Ochieng JB, Omore R, Oundo JO, Hossain A, Das SK, Ahmed S, Qureshi S, Quadri F, Adegbola RA, Antonio M, Hossain MJ, Akinsola A, Mandomando I, Nhampossa T, Acacio S, Biswas K, O’Reilly CE, Mintz ED, Berkeley LY, Muhsen K, Sommerfelt H, Robins-Browne RM, Levine MM. Burden and aetiology of diarrhoeal disease in infants and young children in developing countries (the Global Enteric Multicenter Study, GEMS): a prospective, case-control study. Lancet. 2013;382(9888):209–222. PMID: 23680352.

17. Templeton TJ, Iyer LM, Anantharaman V, Enomoto S, Abrahante JE, Subramanian GM, Hoffman SL, Abrahamsen MS, Aravind L. Comparative analysis of apicomplexa and genomic diversity in eukaryotes. Gen Res. 2004;14:1686–1695.

18. Berney C, Pawlowski J. A molecular time-scale for eukaryote evolution recalibrated with the continuous microfossil record. Proc Biol Sci. 2006;273(1596):1867–1872. PMID: 16822745; 1634798.

19. Li L, Stoeckert CJ, Jr., Roos DS. OrthoMCL: identification of ortholog groups for eukaryotic genomes. Genome Res. 2003;13(9):2178–2189. PMID: 12952885; 403725.

20. Hyde JE. Targeting purine and pyrimidine metabolism in human apicomplexan parasites. Curr Drug Targets. 2007;8(1):31–47. PMID: 17266529; PMC2720675.

21. Neville AJ, Zach SJ, Wang X, Larson JJ, Judge AK, Davis LA, Vennerstrom JL, Davis PH. Clinically Available Medicines Demonstrating Anti-Toxoplasma Activity. Antimicrob Agents Chemother. 2015;59(12):7161–7169. PMID: 26392504; PMC4649158.

22. Wei HX, Wei SS, Lindsay DS, Peng HJ. A Systematic Review and Meta-Analysis of the Efficacy of Anti-Toxoplasma gondii Medicines in Humans. PLoS One. 2015;10(9):e0138204. PMID: 26394212; PMC4578932.

23. McCabe RE. Antitoxoplasma chemotherapy. In: Joynson DHM, Wreghitt TG, editors. Toxoplasmosis: a comprehensive clinical guide. Cambridge: Cambridge Univ. Press; 2001. p. 319–359.

24. Ben-Harari RR, Goodwin E, Casoy J. Adverse Event Profile of Pyrimethamine-Based Therapy in Toxoplasmosis: A Systematic Review. Drugs R D. 2017;17(4):523–544. PMID: 28879584; PMC5694419.

25. Hernandez AV, Thota P, Pellegrino D, Pasupuleti V, Benites-Zapata VA, Deshpande A, Penalva de Oliveira AC, Vidal JE. A systematic review and meta-analysis of the relative efficacy and safety of treatment regimens for HIV-associated cerebral toxoplasmosis: is trimethoprim-sulfamethoxazole a real option? HIV Med. 2017;18(2):115–124. PMID: 27353303.

26. Kieffer F, Wallon M. Congenital toxoplasmosis. Handb Clin Neurol. 2013;112:1099–1101. PMID: 23622316.

27. Adeyemi OS, Sugi T, Han Y, Kato K. Screening of chemical compound libraries identified new anti-Toxoplasma gondii agents. Parasitol Res. 2018;117(2):355–363. PMID: 29260298.

28. Benmerzouga I, Checkley LA, Ferdig MT, Arrizabalaga G, Wek RC, Sullivan WJ, Jr. Guanabenz repurposed as an antiparasitic with activity against acute and latent toxoplasmosis. Antimicrob Agents Chemother. 2015;59(11):6939–6945. PMID: 26303803; PMC4604420.

29. Konrad C, Queener SF, Wek RC, Sullivan WJ, Jr. Inhibitors of eIF2alpha dephosphorylation slow replication and stabilize latency in Toxoplasma gondii. Antimicrob Agents Chemother. 2013;57(4):1815–1822. PMID: 23380722; PMC3623309.

30. Dittmar AJ, Drozda AA, Blader IJ. Drug Repurposing Screening Identifies Novel Compounds That Effectively Inhibit Toxoplasma gondii Growth. mSphere. 2016;1(2). PMID: 27303726; PMC4894684.

31. Eastman RT, Fidock DA. Artemisinin-based combination therapies: a vital tool in efforts to eliminate malaria. Nat Rev Microbiol. 2009;7(12):864–874. PMID: 19881520; PMC2901398.

32. Wellems TE, Plowe CV. Chloroquine-resistant malaria. Journal of Infectious Diseases. 2001;184:770–776.

33. O’Neill PM, Barton VE, Ward SA. The molecular mechanism of action of artemisinin--the debate continues. Molecules. 2010;15(3):1705–1721. PMID: 20336009.

34. Jones-Brando L, D’Angelo J, Posner GH, Yolken RH. *In vitro* inhibition of *Toxoplasma gondii* by four new derivatives of artemisinin. Antimicrob Agents Chem. 2006;50:4206–4208.

35. Nagamune K, Moreno SNJ, Sibley LD. Artemisinin resistant mutants of *Toxoplasma gondii* have altered calcium homeostasis. Anti Microb Agents Chemother. 2007;51:3816–3823.

36. Hencken CP, Jones-Brando L, Bordon C, Stohler R, Mott BT, Yolken R, Posner GH, Woodard LE. Thiazole, oxadiazole, and carboxamide derivatives of artemisinin are highly selective and potent inhibitors of Toxoplasma gondii. J Med Chem. 2010;53(9):3594–3601. PMID: 20373807; PMC2865576.

37. Dunay IR, Chan WC, Haynes RK, Sibley LD. Artemisone and artemiside control acute and reactivated toxoplasmosis in the murine model. Antimicrob Agents Chemother. 2009;53:4450 - 4456.

38. Schultz TL, Hencken CP, Woodard LE, Posner GH, Yolken RH, Jones-Brando L, Carruthers VB. A thiazole derivative of artemisinin moderately reduces Toxoplasma gondii cyst burden in infected mice. J Parasitol. 2014;100(4):516–521. PMID: 24524228.

39. Flannery EL, Chatterjee AK, Winzeler EA. Antimalarial drug discovery - approaches and progress towards new medicines. Nat Rev Microbiol. 2017;15(9):572. PMID: 28736448.

40. Yeung BK, Zou B, Rottmann M, Lakshminarayana SB, Ang SH, Leong SY, Tan J, Wong J, Keller-Maerki S, Fischli C, Goh A, Schmitt EK, Krastel P, Francotte E, Kuhen K, Plouffe D, Henson K, Wagner T, Winzeler EA, Petersen F, Brun R, Dartois V, Diagana TT, Keller TH. Spirotetrahydro beta-carbolines (spiroindolones): a new class of potent and orally efficacious compounds for the treatment of malaria. J Med Chem. 2010;53(14):5155–5164. PMID: 20568778.

41. van Pelt-Koops JC, Pett HE, Graumans W, van der Vegte-Bolmer M, van Gemert GJ, Rottmann M, Yeung BK, Diagana TT, Sauerwein RW. The spiroindolone drug candidate NITD609 potently inhibits gametocytogenesis and blocks Plasmodium falciparum transmission to anopheles mosquito vector. Antimicrob Agents Chemother. 2012;56(7):3544–3548. PMID: 22508309; PMC3393464.

42. Rottmann M, McNamara C, Yeung BK, Lee MC, Zou B, Russell B, Seitz P, Plouffe DM, Dharia NV, Tan J, Cohen SB, Spencer KR, Gonzalez-Paez GE, Lakshminarayana SB, Goh A, Suwanarusk R, Jegla T, Schmitt EK, Beck HP, Brun R, Nosten F, Renia L, Dartois V, Keller TH, Fidock DA, Winzeler EA, Diagana TT. Spiroindolones, a potent compound class for the treatment of malaria. Science. 2010;329(5996):1175–1180. PMID: 20813948; PMC3050001.

43. Spillman NJ, Allen RJ, McNamara CW, Yeung BK, Winzeler EA, Diagana TT, Kirk K. Na(+) regulation in the malaria parasite Plasmodium falciparum involves the cation ATPase PfATP4 and is a target of the spiroindolone antimalarials. Cell Host Microbe. 2013;13(2):227–237. PMID: 23414762; PMC3574224.

44. Van Voorhis WC, Adams JH, Adelfio R, Ahyong V, Akabas MH, Alano P, Alday A, Aleman Resto Y, Alsibaee A, Alzualde A, Andrews KT, Avery SV, Avery VM, Ayong L, Baker M, Baker S, Ben Mamoun C, Bhatia S, Bickle Q, Bounaadja L, Bowling T, Bosch J, Boucher LE, Boyom FF, Brea J, Brennan M, Burton A, Caffrey CR, Camarda G, Carrasquilla M, Carter D, Belen Cassera M, Chih-Chien Cheng K, Chindaudomsate W, Chubb A, Colon BL, Colon-Lopez DD, Corbett Y, Crowther GJ, Cowan N, D’Alessandro S, Le Dang N, Delves M, DeRisi JL, Du AY, Duffy S, Abd El-Salam El-Sayed S, Ferdig MT, Fernandez Robledo JA, Fidock DA, Florent I, Fokou PV, Galstian A, Gamo FJ, Gokool S, Gold B, Golub T, Goldgof GM, Guha R, Guiguemde WA, Gural N, Guy RK, Hansen MA, Hanson KK, Hemphill A, Hooft van Huijsduijnen R, Horii T, Horrocks P, Hughes TB, Huston C, Igarashi I, Ingram-Sieber K, Itoe MA, Jadhav A, Naranuntarat Jensen A, Jensen LT, Jiang RH, Kaiser A, Keiser J, Ketas T, Kicka S, Kim S, Kirk K, Kumar VP, Kyle DE, Lafuente MJ, Landfear S, Lee N, Lee S, Lehane AM, Li F, Little D, Liu L, Llinas M, Loza MI, Lubar A, Lucantoni L, Lucet I, Maes L, Mancama D, Mansour NR, March S, McGowan S, Medina Vera I, Meister S, Mercer L, Mestres J, Mfopa AN, Misra RN, Moon S, Moore JP, Morais Rodrigues da Costa F, Muller J, Muriana A, Nakazawa Hewitt S, Nare B, Nathan C, Narraidoo N, Nawaratna S, Ojo KK, Ortiz D, Panic G, Papadatos G, Parapini S, Patra K, Pham N, Prats S, Plouffe DM, Poulsen SA, Pradhan A, Quevedo C, Quinn RJ, Rice CA, Abdo Rizk M, Ruecker A, St Onge R, Salgado Ferreira R, Samra J, Robinett NG, Schlecht U, Schmitt M, Silva Villela F, Silvestrini F, Sinden R, Smith DA, Soldati T, Spitzmuller A, Stamm SM, Sullivan DJ, Sullivan W, Suresh S, Suzuki BM, Suzuki Y, Swamidass SJ, Taramelli D, Tchokouaha LR, Theron A, Thomas D, Tonissen KF, Townson S, Tripathi AK, Trofimov V, Udenze KO, Ullah I, Vallieres C, Vigil E, Vinetz JM, Voong Vinh P, Vu H, Watanabe NA, Weatherby K, White PM, Wilks AF, Winzeler EA, Wojcik E, Wree M, Wu W, Yokoyama N, Zollo PH, Abla N, Blasco B, Burrows J, Laleu B, Leroy D, Spangenberg T, Wells T, Willis PA. Open Source Drug Discovery with the Malaria Box Compound Collection for Neglected Diseases and Beyond. PLoS Pathog. 2016;12(7):e1005763. PMID: 27467575; PMC4965013.

45. Boyom FF, Fokou PV, Tchokouaha LR, Spangenberg T, Mfopa AN, Kouipou RM, Mbouna CJ, Donfack VF, Zollo PH. Repurposing the open access malaria box to discover potent inhibitors of Toxoplasma gondii and Entamoeba histolytica. Antimicrob Agents Chemother. 2014;58(10):5848–5854. PMID: 25049259; PMC4187973.

46. Doggett JS, Nilsen A, Forquer I, Wegmann KW, Jones-Brando L, Yolken RH, Bordon C, Charman SA, Katneni K, Schultz T, Burrows JN, Hinrichs DJ, Meunier B, Carruthers VB, Riscoe MK. Endochin-like quinolones are highly efficacious against acute and latent experimental toxoplasmosis. Proc Natl Acad Sci U S A. 2012;109(39):15936–15941. PMID: 23019377; 3465437.

47. Kovacs JA. Efficacy of atovoquone in treatment of toxoplasmosis in AIDS patients. Lancet. 1992;340:637–638.

48. Subramanian G, Belekar MA, Shukla A, Tong JX, Sinha A, Chu TTT, Kulkarni AS, Preiser PR, Reddy DS, Tan KSW, Shanmugam D, Chandramohanadas R. Targeted Phenotypic Screening in Plasmodium falciparum and Toxoplasma gondii Reveals Novel Modes of Action of Medicines for Malaria Venture Malaria Box Molecules. mSphere. 2018;3(1). PMID: 29359192; PMC5770543.

49. Spalenka J, Escotte-Binet S, Bakiri A, Hubert J, Renault JH, Velard F, Duchateau S, Aubert D, Huguenin A, Villena I. Discovery of New Inhibitors of Toxoplasma gondii via the Pathogen Box. Antimicrob Agents Chemother. 2018;62(2). PMID: 29133550; PMC5786798.

50. Wells TN, Hooft van Huijsduijnen R, Van Voorhis WC. Malaria medicines: a glass half full? Nat Rev Drug Discov. 2015;14(6):424–442. PMID: 26000721.

51. Chio LC, Queener SF. Identification of highly potent and selective inhibitors of Toxoplasma gondii dihydrofolate reductase. Antimicrob Agents Chemother. 1993;37(9):1914–1923. PMID: 8239605; PMC188092.

52. Meneceur P, Bouldouyre MA, Aubert D, Villena I, Menotti J, Sauvage V, Garin JF, Derouin F. In vitro susceptibility of various genotypic strains of Toxoplasma gondii to pyrimethamine, sulfadiazine, and atovaquone. Antimicrob Agents Chemother. 2008;52(4):1269–1277. PMID: 18212105; 2292506.

53. Harris C, Salgo MP, Tanowitz HB, Wittner M. In vitro assessment of antimicrobial agents against *Toxoplasma gondii*. J Infect Dis. 1988;157:14–22.

54. Stahl W, Collins DN, Benitez P, Turek G, Gaafar H. Toxoplasma gondii: disease patterns in mice treated with the folate antagonist methotrexate. Exp Parasitol. 1976;39(1):135–142. PMID: 1253880.

55. Holfels E, McAuley J, Mack D, Milhous WK, McLeod R. *In vitro* effects of artemisinin ether, cycloguanil hydrochloride (alone and in combination with sulfadiazine), quinine sulfate, mefloquine, primaquine phosphate, trifluoperazine hydrochloride, and verapamil on *Toxoplasma gondii*. Antimicrob Agents Chem. 1994;38:1392–1396.

56. Reynolds MG, Roos DS. A biochemical and genetic model for parasite resistance to antifolates. Toxoplasma gondii provides insights into pyrimethamine and cycloguanil resistance in Plasmodium falciparum. J Biol Chem. 1998;273(6):3461–3469. PMID: 9452469.

57. Belal US, Norose K, Mohamed RM, Sekine S, Nukaga T, Ito K, Abdellatif MZ, Abdelgelil NH, Yano A. Quantitative assessment of the effects of sulfamethoxazole on Toxoplasma gondii loads in susceptible WT C57BL/6 mice as an immunocompetent host model. Parasitol Int. 2016;65(1):1–4. PMID: 26384856.

58. Brun-Pascaud M, Chau F, Garry L, Jacobus D, Derouin F, Girard PM. Combination of PS-15, epiroprim, or pyrimethamine with dapsone in prophylaxis of Toxoplasma gondii and Pneumocystis carinii dual infection in a rat model. Antimicrob Agents Chemother. 1996;40(9):2067–2070. PMID: 8878582; PMC163474.

59. Derouin F, Almadany R, Chau F, Rouveix B, Pocidalo JJ. Synergistic activity of azithromycin and pyrimethamine or sulfadiazine in acute experimental toxoplasmosis. Antimicrob Agents Chemother. 1992;36(5):997–1001. PMID: 1324642; PMC188824.

60. Chang HR, Comte R, Pechére JC. In vitro and in vivo efects of doxycycline on *Toxoplasma gondii*. Anti Microb Agents Chemother. 1990;34:775–780.

61. Berg-Candolfi M, Candolfi E. Depression of the N-demethylation of erythromycin, azithromycin, clarithromycin and clindamycin in murine Toxoplasma infection. Int J Parasitol. 1996;26(11):1321–1323. PMID: 9024879.

62. Djurkovic-Djakovic O, Milenkovic V, Nikolic A, Bobic B, Grujic J. Efficacy of atovaquone combined with clindamycin against murine infection with a cystogenic (Me49) strain of Toxoplasma gondii. J Antimicrob Chemother. 2002;50(6):981–987. PMID: 12461021.

63. Djurkovic-Djakovic O, Nikolic T, Robert-Gangneux F, Bobic B, Nikolic A. Synergistic effect of clindamycin and atovaquone in acute murine toxoplasmosis. Antimicrob Agents Chemother. 1999;43(9):2240–2244. PMID: 10471572; PMC89454.

64. Ribeiro M, Franco PS, Lopes-Maria JB, Angeloni MB, Barbosa BF, Gomes AO, Castro AS, Silva RJD, Oliveira FC, Milian ICB, Martins-Filho OA, Ietta F, Mineo JR, Ferro EAV. Azithromycin treatment is able to control the infection by two genotypes of Toxoplasma gondii in human trophoblast BeWo cells. Exp Parasitol. 2017;181:111–118. PMID: 28803905.

65. Camps M, Arrizabalaga G, Boothroyd J. An rRNA mutation identifies the apicoplast as the target for clindamycin in Toxoplasma gondii. Mol Microbiol. 2002;43(5):1309–1318. PMID: 11918815.

66. Slimings C, Riley TV. Antibiotics and hospital-acquired Clostridium difficile infection: update of systematic review and meta-analysis. J Antimicrob Chemother. 2014;69(4):881–891. PMID: 24324224.

67. Alday PH, Bruzual I, Nilsen A, Pou S, Winter R, Ben Mamoun C, Riscoe MK, Doggett JS. Genetic Evidence for Cytochrome b Qi Site Inhibition by 4(1H)-Quinolone-3-Diarylethers and Antimycin in Toxoplasma gondii. Antimicrob Agents Chemother. 2017;61(2). PMID: 27919897; PMC5278733.

68. McFadden DC, Tomavo S, Berry EA, Boothroyd JC. Characterization of cytochrome b from *Toxoplasma gondii* and Qo domain mutations as a mechanism of atovaquone^-^ resistance. Mol Biochem Parasitol. 2000;108(1):1–12.

69. Dunay IR, Heimesaat MM, Bushrab FN, Muller RH, Stocker H, Arasteh K, Kurowski M, Fitzner R, Borner K, Liesenfeld O. Atovaquone maintenance therapy prevents reactivation of toxopasmic encepahalitis in the murine model of reactivated toxoplasmosis. Antimicrob Agents Chem. 2004;48:4848–4854.

70. Gajurel K, Gomez CA, Dhakal R, Vogel H, Montoya JG. Failure of primary atovaquone prophylaxis for prevention of toxoplasmosis in hematopoietic cell transplant recipients. Transpl Infect Dis. 2016;18(3):446–452. PMID: 27016655.

71. McPhillie M, Zhou Y, El Bissati K, Dubey J, Lorenzi H, Capper M, Lukens AK, Hickman M, Muench S, Verma SK, Weber CR, Wheeler K, Gordon J, Sanders J, Moulton H, Wang K, Kim TK, He Y, Santos T, Woods S, Lee P, Donkin D, Kim E, Fraczek L, Lykins J, Esaa F, Alibana-Clouser F, Dovgin S, Weiss L, Brasseur G, Wirth D, Kent M, Hood L, Meunieur B, Roberts CW, Hasnain SS, Antonyuk SV, Fishwick C, McLeod R. New paradigms for understanding and step changes in treating active and chronic, persistent apicomplexan infections. Sci Rep. 2016;6:29179. PMID: 27412848; PMC4944145.

72. Frueh L, Li Y, Mather MW, Li Q, Pou S, Nilsen A, Winter RW, Forquer IP, Pershing AM, Xie LH, Smilkstein MJ, Caridha D, Koop DR, Campbell RF, Sciotti RJ, Kreishman-Deitrick M, Kelly JX, Vesely B, Vaidya AB, Riscoe MK. Alkoxycarbonate Ester Prodrugs of Preclinical Drug Candidate ELQ-300 for Prophylaxis and Treatment of Malaria. ACS Infect Dis. 2017;3(10):728–735. PMID: 28927276.

73. Sullivan DJ, Jr., Matile H, Ridley RG, Goldberg DE. A common mechanism for blockade of heme polymerization by antimalarial quinolines. J Biol Chem. 1998;273(47):31103–31107. PMID: 9813011.

74. Tekwani BL, Walker LA. 8-Aminoquinolines: future role as antiprotozoal drugs. Curr Opin Infect Dis. 2006;19(6):623–631. PMID: 17075340.

75. Rudrapal M, Chetia D. Endoperoxide antimalarials: development, structural diversity and pharmacodynamic aspects with reference to 1,2,4-trioxane-based structural scaffold. Drug Des Devel Ther. 2016;10:3575–3590. PMID: 27843298; PMC5098533.

76. Spillman NJ, Kirk K. The malaria parasite cation ATPase PfATP4 and its role in the mechanism of action of a new arsenal of antimalarial drugs. Int J Parasitol Drugs Drug Resist. 2015;5(3):149–162. PMID: 26401486; PMC4559606.

77. Zhou Y, Fomovska A, Muench S, Lai BS, Mui E, McLeod R. Spiroindolone that inhibits PfATPase4 is a potent, cidal inhibitor of Toxoplasma gondii tachyzoites in vitro and in vivo. Antimicrob Agents Chemother. 2014;58(3):1789–1792. PMID: 24366743; PMC3957861.

78. Vaidya AB, Morrisey JM, Zhang Z, Das S, Daly TM, Otto TD, Spillman NJ, Wyvratt M, Siegl P, Marfurt J, Wirjanata G, Sebayang BF, Price RN, Chatterjee A, Nagle A, Stasiak M, Charman SA, Angulo-Barturen I, Ferrer S, Belen Jimenez-Diaz M, Martinez MS, Gamo FJ, Avery VM, Ruecker A, Delves M, Kirk K, Berriman M, Kortagere S, Burrows J, Fan E, Bergman LW. Pyrazoleamide compounds are potent antimalarials that target Na+ homeostasis in intraerythrocytic Plasmodium falciparum. Nature communications. 2014;5:5521. PMID: 25422853; PMC4263321.

79. White NJ, Duong TT, Uthaisin C, Nosten F, Phyo AP, Hanboonkunupakarn B, Pukrittayakamee S, Jittamala P, Chuthasmit K, Cheung MS, Feng Y, Li R, Magnusson B, Sultan M, Wieser D, Xun X, Zhao R, Diagana TT, Pertel P, Leong FJ. Antimalarial Activity of KAF156 in Falciparum and Vivax Malaria. N Engl J Med. 2016;375(12):1152–1160. PMID: 27653565; PMC5142602.

80. Dembele L, Ang X, Chavchich M, Bonamy GMC, Selva JJ, Lim MY, Bodenreider C, Yeung BKS, Nosten F, Russell BM, Edstein MD, Straimer J, Fidock DA, Diagana TT, Bifani P. The Plasmodium PI(4)K inhibitor KDU691 selectively inhibits dihydroartemisinin-pretreated Plasmodium falciparum ring-stage parasites. Sci Rep. 2017;7(1):2325. PMID: 28539634; PMC5443816.

81. Hoepfner D, McNamara CW, Lim CS, Studer C, Riedl R, Aust T, McCormack SL, Plouffe DM, Meister S, Schuierer S, Plikat U, Hartmann N, Staedtler F, Cotesta S, Schmitt EK, Petersen F, Supek F, Glynne RJ, Tallarico JA, Porter JA, Fishman MC, Bodenreider C, Diagana TT, Movva NR, Winzeler EA. Selective and specific inhibition of the plasmodium falciparum lysyl-tRNA synthetase by the fungal secondary metabolite cladosporin. Cell Host Microbe. 2012;11(6):654–663. PMID: 22704625; PMC3391680.

82. Khan S, Sharma A, Belrhali H, Yogavel M, Sharma A. Structural basis of malaria parasite lysyl-tRNA synthetase inhibition by cladosporin. J Struct Funct Genomics. 2014;15(2):63–71. PMID: 24935905.

83. Phillips MA, Rathod PK. Plasmodium dihydroorotate dehydrogenase: a promising target for novel anti-malarial chemotherapy. Infect Disord Drug Targets. 2010;10(3):226–239. PMID: 20334617; PMC2883174.

84. Phillips MA, Lotharius J, Marsh K, White J, Dayan A, White KL, Njoroge JW, El Mazouni F, Lao Y, Kokkonda S, Tomchick DR, Deng X, Laird T, Bhatia SN, March S, Ng CL, Fidock DA, Wittlin S, Lafuente-Monasterio M, Benito FJ, Alonso LM, Martinez MS, Jimenez-Diaz MB, Bazaga SF, Angulo-Barturen I, Haselden JN, Louttit J, Cui Y, Sridhar A, Zeeman AM, Kocken C, Sauerwein R, Dechering K, Avery VM, Duffy S, Delves M, Sinden R, Ruecker A, Wickham KS, Rochford R, Gahagen J, Iyer L, Riccio E, Mirsalis J, Bathhurst I, Rueckle T, Ding X, Campo B, Leroy D, Rogers MJ, Rathod PK, Burrows JN, Charman SA. A long-duration dihydroorotate dehydrogenase inhibitor (DSM265) for prevention and treatment of malaria. Sci Transl Med. 2015;7(296):296ra111. PMID: 26180101; PMC4539048.

85. Hortua Triana MA, Cajiao Herrera D, Zimmermann BH, Fox BA, Bzik DJ. Pyrimidine Pathway-Dependent and -Independent Functions of the Toxoplasma gondii Mitochondrial Dihydroorotate Dehydrogenase. Infect Immun. 2016;84(10):2974–2981. PMID: 27481247; PMC5038078.

86. Pfefferkorn ER, Pfefferkorn LC. Specific labeling of intracellular *Toxoplasma gondii* with uracil. Journal of Protozoology. 1977;24:449–453.

87. Tobin CM, Knoll LJ. A patatin-like protein protects Toxoplasma gondii from degradation in a nitric oxide-dependent manner. Infect Immun. 2012;80(1):55–61. PMID: 22006568; PMC3255658.

